# A single amino acid substitution in the Borna disease virus glycoprotein enhances the infectivity titer of vesicular stomatitis virus pseudotyped virus by altering membrane fusion activity

**DOI:** 10.1101/2024.05.22.595257

**Authors:** Yusa Akiba, Hiromichi Matsugo, Takehiro Kanda, Modoka Sakai, Akiko Makino, Keizo Tomonaga

**Affiliations:** Laboratory of RNA Viruses, Department of Virus Research, Institute for Life and Medical Sciences, Kyoto University; Department of Mammalian Regulatory Network, Graduate School of Biostudies, Kyoto University; Department of Molecular Virology, Graduate School of Medicine, Kyoto University

**Keywords:** Borna disease virus, cell entry, membrane fusion, pseudotyped virus, vesicular stomatitis virus

## Abstract

Borna disease virus 1 (BoDV-1) causes acute fatal encephalitis in mammals, including humans. Despite its importance, research on BoDV-1 cell entry has been hindered by low viral particle production in cells and the lack of cytopathic effects. To address these issues, we developed a method to efficiently produce vesicular stomatitis virus (VSV) pseudotyped with glycoprotein (G) of the genus *Orthobornavirus*, including BoDV-1. We discovered that optimal G expression is required to obtain a high infectivity titer of the VSV pseudotyped virus. Remarkably, the infectivity of the VSV pseudotyped virus with G from the BoDV-1 strain huP2br was significantly higher than that of the VSV pseudotyped virus with G from the He/80 strain. Mutational analysis demonstrated that the BoDV-1-G residue 307 determines the infectivity titer of VSV pseudotyped with BoDV-1-G (VSV-BoDV-1-G). A cell‒cell fusion assay indicated that this residue plays a pivotal role in membrane fusion, thus suggesting that high membrane fusion activity and a broad pH range for membrane fusion are crucial for achieving a high infectivity titer of VSV-BoDV-1-G. This finding may be extended to increase the infectivity titer of VSV pseudotyped virus with other orthobornavirus G. Our study also contributes to identifying functional domains of BoDV-1-G and provides insight into G-mediated cell entry.

## Introduction

Borna disease virus 1 (BoDV-1) is a nonsegmented, negative-strand RNA virus that belongs to the genus *Orthobornavirus* in the family *Bornaviridae*. The bicolored white-toothed shrew (*Crocidura leucodon*) is considered the indigenous host of BoDV-1. BoDV-1 causes fatal encephalitis in various mammals, including cattle, horses, and sheep^1^. Although its association with mental disorders in humans has remained controversial for decades due to its neurotropic property and ability to establish persistent infection^2–5^, this association is now questionable. In 2015, a related virus known as variegated squirrel bornavirus 1 (VSBV-1) was identified as the etiologic agent for fatal encephalitis in humans^6^. Subsequent retrospective studies demonstrated that BoDV-1 also causes fatal encephalitis in humans^7,8^. On the other hand, avian bornaviruses (ABV), which also belong to the genus *Orthobornavirus*, are known to cause proventricular dilatation disease (PDD) in birds^9^.

The genome of BoDV-1 is approximately 8.9 kb and encodes at least six proteins: nucleoprotein (N), phosphoprotein (P), X protein, matrix protein (M), glycoprotein (G), and RNA-dependent RNA polymerase (L). N, P, and L are essential for the formation of viral ribonucleoproteins (vRNPs), transcription, and replication of the viral genome. M is involved in particle formation, whereas G is essential for cell entry and cell-to-cell spread^10,11^.

G is initially synthesized as precursor G (Pre-G), which is cleaved into GP1 and GP2 by furin and undergoes glycosylation^12,13^. BoDV-1 binds to receptors via GP1 and is internalized by endocytosis^14–16^. Subsequently, endosomal acidification induces a conformational change in G, thus leading to membrane fusion and the release of vRNPs into the cytoplasm^17^. Although some host factors involved in cell entry have been reported, the entry mechanism remains largely unknown^18,19^.

BoDV-1 produces few viral particles within infected cells and is less likely to release viral particles outside of infected cells^20,21^. Therefore, considerable effort and specialized equipment are required to obtain high titers of BoDV-1. Moreover, because BoDV-1 does not cause cytopathic effects in infected cells, an indirect fluorescent assay method or recombinant BoDV-1 expressing a reporter gene is required to evaluate BoDV-1 infection. These unique properties of BoDV-1 complicate the analysis of its cell entry mechanism. To overcome these challenges, a convenient system for evaluating cell entry is desired, and we focused on vesicular stomatitis virus (VSV) pseudotyped virus. VSV efficiently incorporates envelope proteins of other viruses into its particles and releases viral particles outside of cells^22^. Therefore, VSV pseudotyped virus with BoDV-1-G (VSV-BoDV-1) expressing a reporter gene serves as a valuable tool for analyzing the cell entry mechanism of BoDV-1. Although VSV-BoDV-1 has been reported, its infectivity titer was relatively low, emphasizing the need for improvement in its production^16^. Additionally, while VSBV-1 is highly pathogenic and requires high biosafety level facilities to handle, replication-deficient VSV pseudotyped virus with VSBV-1-G (VSV-VSBV-1) can be handled at lower biosafety level facilities, thus facilitating the analysis of its cell entry mechanism. Nevertheless, there have been few reports of VSV pseudotyped virus with orthobornavirus G other than BoDV-1-G.

To address these problems, we produced VSV pseudotyped virus with various orthobornavirus G, including BoDV-1-G, and found that optimizing the G expression level is critical for efficient production. The infectivity of the VSV pseudotyped virus with BoDV-1-huP2br-G was approximately 200-fold higher than that of BoDV-1-He/80-G, and this difference was attributed to a variation at BoDV-1-G residue 307. Furthermore, this residue affected membrane fusion activity and the pH threshold for membrane fusion. Our findings indicate that membrane fusion activity and the pH threshold for membrane fusion of G are crucial for increasing the infectivity of VSV-BoDV-1. This insight can be extended to enhance the infectivity titer of VSV pseudotyped virus with other orthobornavirus G, including those that may emerge in the future.

## Materials and methods

### Cells

Human embryonic kidney (293T) cells and African green monkey kidney (Vero) cells were cultured in Dulbecco’s modified Eagle medium (DMEM; Thermo Fisher Scientific, Waltham, MA, USA) supplemented with 10% fetal calf serum (FCS) and 1% penicillin/streptomycin (Nacalai Tesque, Kyoto, Japan). Human oligodendroglioma (OL) cells were cultured in DMEM supplemented with 5% FCS and 1% penicillin/streptomycin. The cells were maintained at 37 °C with 5% CO_2_.

### Plasmids

To construct plasmids expressing BoDV-1-G, the G genes of the He/80 (GenBank accession no. AJ311522) and huP2br (GenBank accession no. AB258389) strains were amplified, and its splicing donor and splicing acceptor sites were mutated using site-directed mutagenesis PCR, as described previously^13^. To construct a plasmid expressing BoDV-2-G, the G gene of BoDV-2 (GenBank accession no. AJ311524) was amplified using PCR. These PCR products were inserted into EcoRI-XhoI-digested linearized pCAGGS using the NEBuilder HiFi DNA Assembly Cloning Kit (New England Biolabs, Ipswich, MA, USA). Previously constructed plasmids expressing G of VSBV-1, canary bornavirus 1 (CnBV-1), parrot bornavirus-2 (PaBV-2), PaBV-4, and PaBV-5 were also used^13^. Plasmids expressing a series of single amino acid-substituted BoDV-1-G were constructed using site-directed mutagenesis PCR.

### Western blotting

To prepare protein samples, cells and ultracentrifuged virus pellets were lysed with 2× Laemmli SDS sample buffer and subsequently boiled at 95 °C for 10 min. Protein samples were subjected to SDS‒PAGE on an e‒PAGE gel (ATTO Corporation, Tokyo, Japan). Afterwards, the proteins were transferred to PVDF membranes using Trans-Blot Turbo PVDF Transfer Packs (Bio-Rad, Hercules, CA, USA), and the membranes were blocked with Blocking One (Nacalai Tesque). After washing the membranes once with TBS-T (Tris-buffered saline containing 0.1% Tween 20), they were incubated with primary antibodies and secondary antibodies using an iBind Flex Western Device (Thermo Fisher Scientific). The following antibodies were utilized in this study: rabbit anti-BoDV-1-G, mouse anti-VSV-M (Merck, Darmstadt, Germany), mouse anti-β-actin (Merck), HRP-conjugated donkey anti-mouse IgG (Jackson ImmunoResearch, West Grove, PA, USA), and HRP-conjugated donkey anti-rabbit IgG (Jackson ImmunoResearch). After washing the membranes twice with ultrapure water, Clarity Western ECL substrate (Bio-Rad) was used for detection.

### Production of the VSV pseudotyped virus

VSV pseudotyped viruses with orthobornavirus G were produced using VSVΔG*-GFP (kindly gifted by Dr. Hideki Tani, Department of Virology, Toyama Institute of Health, Japan), which contains the green fluorescence protein (GFP) gene instead of its own G gene, as previously described with some modifications^23^. 293T cells (1.5×10^6^ cells) seeded onto collagen-coated 6-well plates were transfected with 2.0, 0.2, 0.02 or 0.01 µg of G expression plasmids using PEI Max (Polysciences, Warrington, PA, USA) according to the manufacturer’s instructions. At 24 hours posttransfection, the cells were infected with VSVΔG*-GFP at a multiplicity of infection of 0.6 for 1 hour at 37 °C. The cells were washed twice and incubated with growth medium for 24 hours. Afterwards, the supernatant was collected via centrifugation (5000 × g for 5 min). The virus solutions were stored at -80 °C.

### Titration of the VSV pseudotyped virus

To neutralize the remaining VSVΔG*-GFP, the VSV pseudotyped virus was incubated with the anti-VSV-G neutralizing I1 (kindly gifted by Dr. Masayuki Shimojima, National Institute of Infectious Disease, Japan) for 30 min at 24 °C. Subsequently, Vero cells seeded on 96-well plates were inoculated with 10-fold serial dilutions of neutralized VSV pseudotyped virus for 1 hour at 37 °C. At 24 hours postinfection, GFP-expressing cells were counted as virus-infected cells using a fluorescence microscope (Nikon, Tokyo, Japan).

### Concentration and purification of the VSV pseudotyped virus

The virus solutions were centrifuged at 106800 × g for 3 hours at 4 °C using a SW32Ti rotor (Beckman Coulter, Fullerton, CA, USA). The virus pellets were suspended in phosphate-buffered saline (PBS). They were layered onto a stepwise sucrose gradient (60%, 50%, 30%, and 20%) and centrifuged at 106800 × g for 3 hours at 4 °C. After centrifugation, the virus-containing fraction was collected and centrifuged at 106800 × g for 3 hours at 4 °C.

The virus pellets were subjected to Western blotting analysis.

### Cell‒cell fusion assay

A cell‒cell fusion assay was performed as previously described with some modifications^24,25^. OL cells (3.0×10^5^ cells) seeded onto 12-well plates were cotransfected with G expression plasmids and a GFP expression plasmid. At 24 hours posttransfection, the cells were washed once with PBS containing Mg^2+^ and Ca^2+^ (PBS+) and then incubated with PBS+ adjusted to various pH levels for 5 min at 24 °C. After incubation, the cells were washed with PBS+ twice and incubated with growth medium for 90 min at 37 °C. Fused cells were observed using a fluorescence microscope (Nikon).

### Statistical analysis

The data were statistically analyzed using GraphPad Prism 10 software, and the utilized tests are specified in the figure legends.

## Results

### The infectivity titers of VSV pseudotyped viruses with orthobornavirus G depend on both the strain and the G expression level

Although a previous report showed that VSV pseudotyped virus with BoDV-1-He/80-G could be rescued, reports on pseudotyping with other orthobornavirus G are scarce^16^. To address this, 293T cells were transfected with plasmids expressing various orthobornavirus G and subsequently infected with VSVΔG*-GFP, after which the VSV pseudotyped viruses were harvested. The infectivity titers of the VSV pseudotyped viruses with G of BoDV-1, PaBV-2, and PaBV-4 (VSV-BoDV-1, VSV-PaBV-2, and VSV-PaBV-4) were low but higher than those of the pseudotyped viruses without any envelope protein (VSV-empty), thus indicating successful rescue (Fig. 1A). In contrast, the infectivity titers of VSV pseudotyped viruses with G of BoDV-2, VSBV-1, CnBV-1, and PaBV-5 (VSV-BoDV-2, VSV-VSBV-1, VSV-CnBV-1, and VSV-PaBV-5) were 100-to 2000-fold higher than those of VSV-BoDV-1 (Fig. 1A), highlighting the impact of G on the viral titer.

**Fig. 1.**
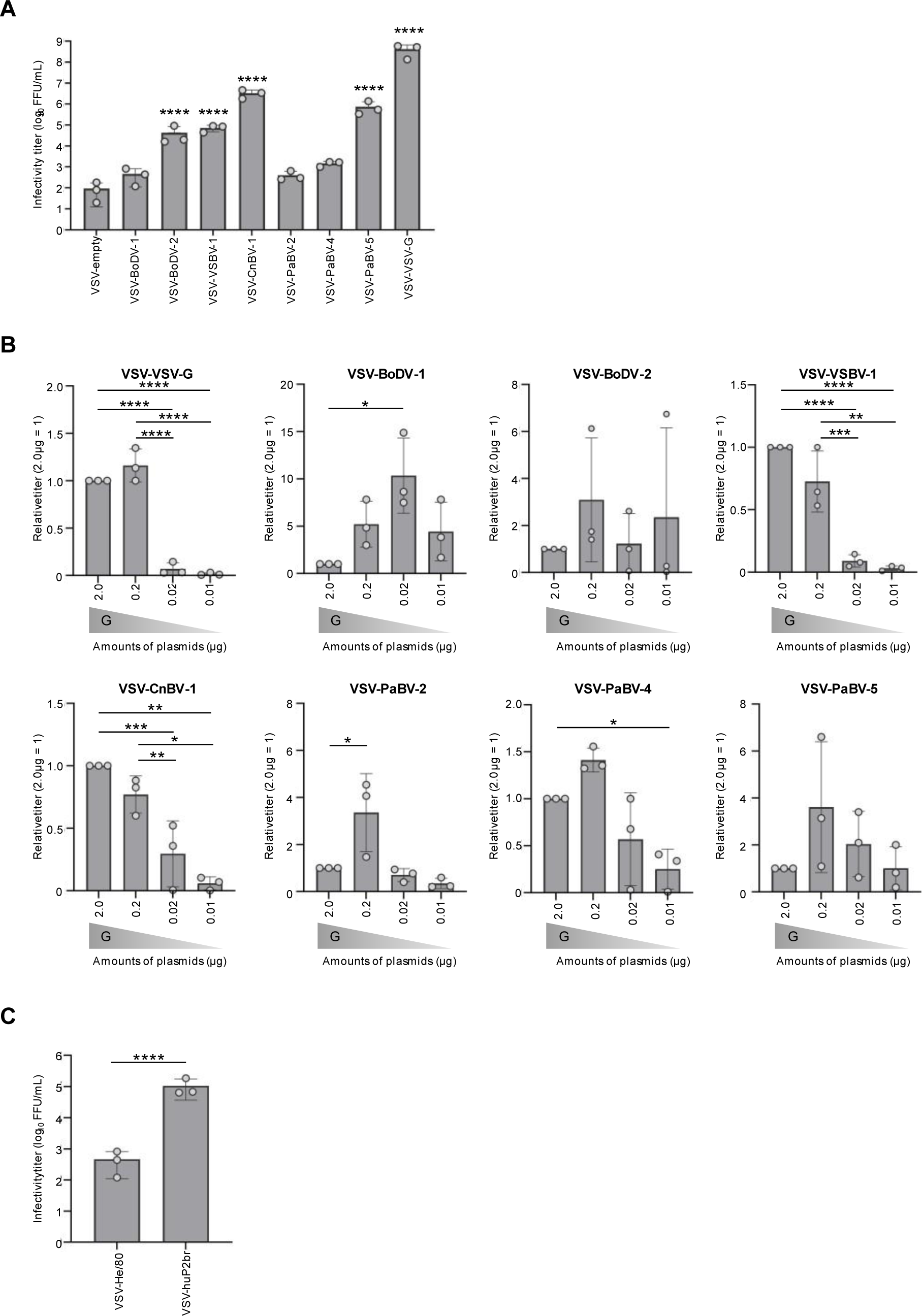
Production of VSV pseudotyped viruses with orthobornavirus G. (A) The infectivity titer of VSV pseudotyped viruses with orthobornavirus G. 293T cells (1.5×10^6^ cells) were transfected with 2.0 µg of G expression plasmids, and the cells were infected with VSVΔG*-GFP. After 24 hours, VSV pseudotyped viruses were harvested. VSV-empty, VSV-BoDV-2, VSV-VSBV-1, VSV-CnBV-1, VSV-PaBV-2, VSV-PaBV-4, and VSV-PaBV-5 were produced simultaneously, whereas VSV-BoDV-1 and VSV-VSV-G were produced at a different time. (B) Comparison of the infectivity titers of VSV pseudotyped viruses produced by various amounts of G expression plasmids. 293T cells (1.5×10^6^ cells) were transfected with 2.0, 0.2, 0.02 or 0.01 µg of G expression plasmids, and the cells were infected with VSVΔG*-GFP. After 24 hours, VSV pseudotyped viruses were harvested. (C) Comparison of the infectivity titers of VSV-He/80 and VSV-huP2br. 293T cells (1.5×10^6^ cells) were transfected with 2.0 µg of plasmids expressing BoDV-1-He/80-G or BoDV-1-huP2br-G, and the cells were infected with VSVΔG*-GFP. After 24 hours, VSV pseudotyped viruses were harvested. The infectivity titer was determined using Vero cells. The bars show the means ± the standard errors (SEs) of three independent experiments. Statistical analysis was performed using one-way ANOVA with Dunnett’s multiple comparisons test (A), with Tukey’s multiple comparisons test (B), or with an unpaired Student’s *t* test (C). *, P < 0.05; **, P < 0.01; ***, P < 0.001; ****, P < 0.0001.

Furin cleavage and glycosylation are key posttranslational modifications of BoDV-1-G. A previous study demonstrated that excess expression of BoDV-1-G leads to impaired proper G cleavage and aberrant glycosylation^13^. To investigate whether optimizing the G expression level could increase the infectivity titer of the VSV pseudotyped virus, 293T cells were transfected with various amounts of G expression plasmids, and the VSV pseudotyped viruses were harvested. Although reducing the G expression did not increase the infectivity titer of the VSV pseudotyped virus with VSV-G, it significantly increased the infectivity titers of VSV-BoDV-1, VSV-BoDV-2, VSV-PaBV-2, VSV-PaBV-4, and VSV-PaBV-5 (Fig. 1B), thus indicating the critical role of optimal G expression in achieving high titers of VSV pseudotyped viruses with orthobornavirus G.

Given the lower infectivity titer of VSV pseudotyped viruses with BoDV-1-G compared to those pseudotyped with other orthobornavirus G, we explored the infectivity titer of VSV pseudotyped virus with G of a different BoDV-1 strain, huP2br. Remarkably, the infectivity titer of the VSV pseudotyped virus with BoDV-1-huP2br-G (VSV-huP2br) was approximately 200-fold higher than that with BoDV-1-He/80-G (VSV-He/80) (Fig. 1C). This discovery highlights that the infectivity titer of VSV pseudotyped viruses varies depending on the viral strain from which G is derived.

### Identification of residues responsible for the difference between VSV-He/80 and VSV-huP2br

Because no studies have compared the G characteristics of different BoDV-1 strains, the mechanism underlying the significant difference in infectivity titer between VSV-He/80 and VSV-huP2br remains unknown. To elucidate this mechanism, we first investigated which residues determine the difference in infectivity titer. We identified ten different residues in G between He/80 and huP2br (Fig. 2A). We constructed a series of a single amino acid substituted G expression plasmids and confirmed their similar expression levels in cells (Fig. 2B and C). Furthermore, we produced VSV pseudotyped viruses with these recombinant G to assess their effect on the viral infectivity titer (Fig. 2D and E). The infectivity titer of VSV-huP2br with the M307I mutation (VSV-huP2br M307I) was approximately 100-fold lower than that of VSV-huP2br wild-type (VSV-huP2br WT) (Fig. 2D). The infectivity titers of VSV-huP2br P56S and VSV-huP2br V185I were also significantly lower than that of VSV-huP2br WT (Fig. 2D). Conversely, the infectivity of VSV-He/80 I307M was 10-fold higher than that of VSV-He/80 WT (Fig. 2E). These results demonstrated that the BoDV-1-G residue 307 is mainly responsible for the difference in the infectivity titer between VSV-He/80 and VSV-huP2br.

**Fig. 2.**
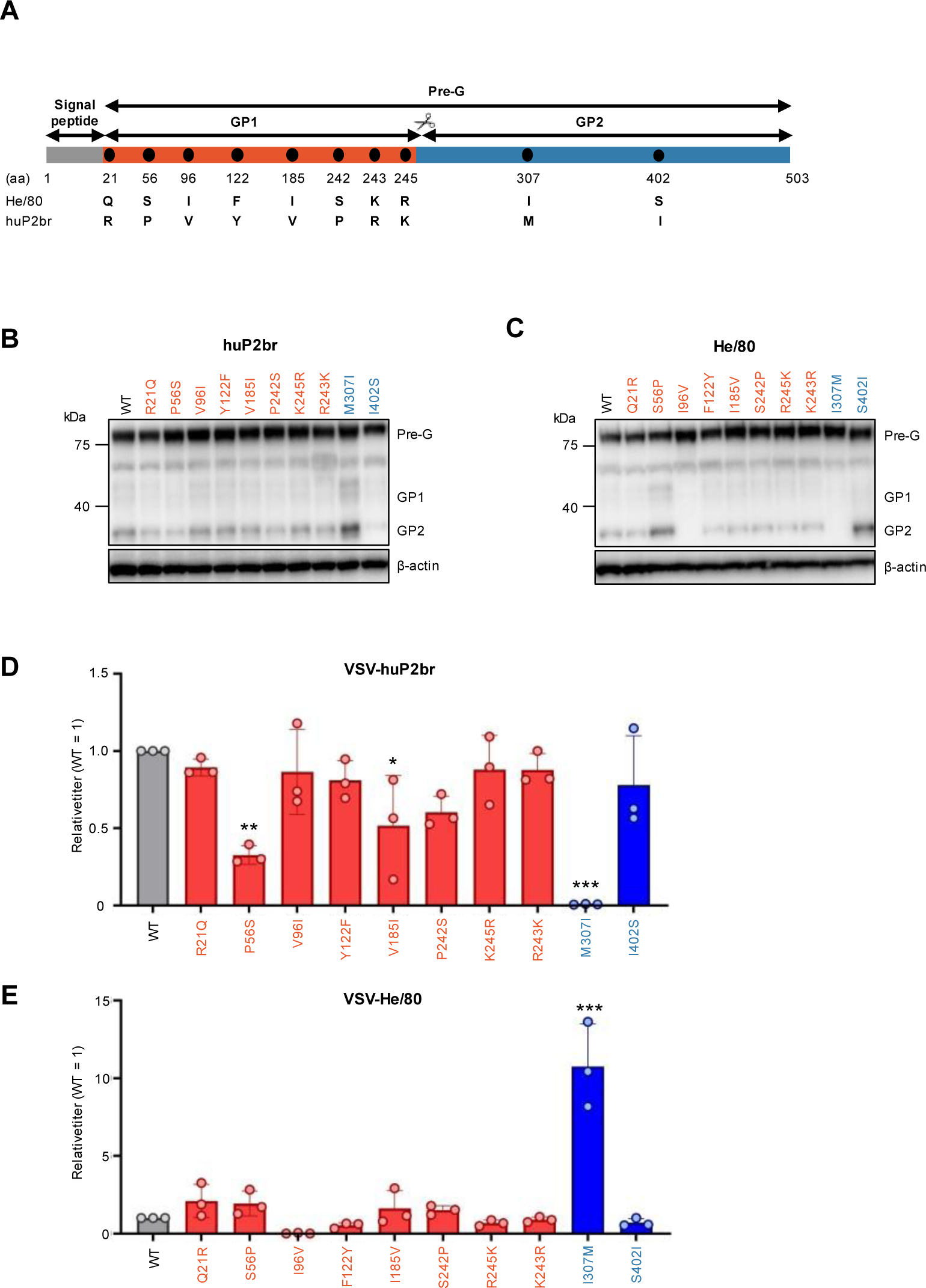
Identification of residues responsible for the difference between VSV-He/80 and VSV-huP2br. (A) Sequence differences in BoDV-1-G between He/80 and huP2br. (B and C) Western blot analysis of G expression in 293T cells at 24 hours posttransfection. Whole lysates of 293T cells transfected with plasmids expressing WT or mutant G of BoDV-1-huP2br (B) or BoDV-1-He/80 (C) were used. (D and E) The infectivity titer of VSV pseudotyped viruses with WT or mutant BoDV-1-huP2br-G (D) or BoDV-1-He/80-G (E). 293T cells (1.5×10^6^ cells) were transfected with 0.02 µg of plasmids expressing BoDV-1-G, and the cells were infected with VSVΔG*-GFP. After 24 hours, VSV pseudotyped viruses were harvested. The infectivity titer was determined using Vero cells. The bars show the means ± the SEs of three independent experiments. Statistical analysis was performed using one-way ANOVA with Dunnett’s multiple comparisons test. *, P < 0.05; **, P < 0.01; ***, P < 0.0001.

### BoDV-1-G residue 307 affects membrane fusion activity and the pH threshold for membrane fusion

Previous research has shown that certain mutations in envelope proteins of other viruses can increase their incorporation into VSV particles, thereby increasing infectivity titers^26–28^. Thus, we investigated whether the BoDV-1-G residue 307 affects its incorporation into VSV particles. We demonstrated that the production of VSV particles, indicated by the presence of VSV-M, was consistent across all of the samples, thus suggesting that VSV budding does not depend on G (Fig. 3A). huP2br WT G was incorporated into VSV particles more efficiently than He/80 WT G (Fig. 3A). Contrary to our expectations, the He/80-G I307M mutation reduced G incorporation into VSV particles (Fig. 3A). Similarly, the huP2br-G M307I mutation slightly decreased G incorporation into VSV particles (Fig. 3A). This observation implies that changes in viral titer due to these mutations occur via mechanisms other than altered G incorporation into VSV particles.

**Fig. 3.**
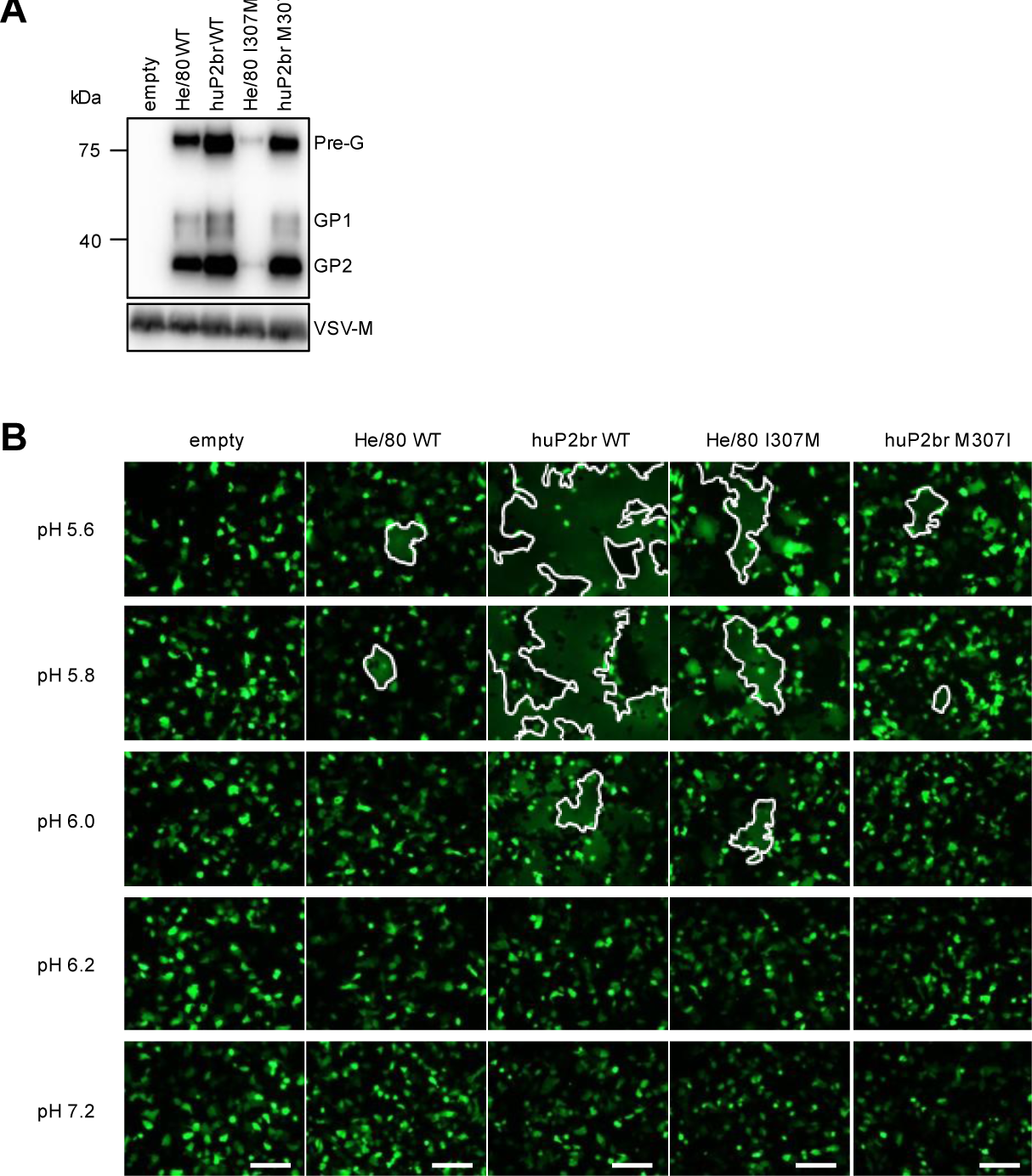
Analysis of the effect of BoDV-1-G residue 307 on the incorporation into VSV particles and the membrane fusion activity of BoDV-1-G. (A) Western blot analysis of VSV-M and BoDV-1-G expression in VSV pseudotyped viruses. Lysates from ultracentrifuged virus pellets were used. (B) Syncytium formation in G-expressing OL cells. OL cells were cotransfected with G expression plasmids and a GFP expression plasmid. At 24 hours posttransfection, the cells were treated with several low-pH buffers. The areas enclosed by the white line indicate syncytia. Scale bar, 20 µm.

Previous reports have shown that membrane fusion activity and the pH threshold for membrane fusion affect the infectivity and replication of other viruses^25,29^. BoDV-1-G residue 307 is located in the GP2 region associated with membrane fusion, thus suggesting that it may affect membrane fusion activity. To test this possibility, we performed a cell‒cell fusion assay. OL cells were cotransfected with G expression plasmids and a GFP expression plasmid. At 24 hours posttransfection, the cells were treated with several low-pH buffers to induce a conformational change in G and syncytium formation. At pH 7.2, no syncytium formation was observed in any G-expressing OL cells (Fig. 3B). At pH 5.6, significant syncytium formation was evident in any G-expressing OL cells (Fig. 3B). The syncytia formed in OL cells expressing He/80 I307M mutant G were larger than those in cells expressing He/80 WT G, whereas those in cells expressing huP2br M307I mutant G were smaller than those in cells expressing huP2br WT G (Fig. 3B). At pH 5.8 and 6.0, limited syncytium formation was observed in cells expressing He/80 WT or huP2br M307I mutant G, whereas significant syncytium formation was observed in cells expressing huP2br WT or He/80 I307M mutant G (Fig. 3B). These results demonstrate that BoDV-1-G residue 307 affects membrane fusion activity and the pH threshold for membrane fusion, thus suggesting that VSV-huP2br can efficiently fuse and infect endosomes within a broader pH range than can VSV-He/80, which subsequently leads to a greater infectivity titer.

## Discussion

The VSV pseudotyped virus system is a powerful tool for the analysis of cell entry mechanisms and for diagnostics. There is a need to develop an efficient method for producing VSV pseudotyped viruses with orthobornavirus G, which has been driven by recent reports of fatal encephalitis in humans caused by BoDV-1 and VSBV-1, along with the global spread of ABV. In this study, we aimed to produce VSV pseudotyped viruses with orthobornavirus G and to elucidate the factors affecting their infectivity titers. We discovered that optimizing the G expression level increased the infectivity titer of the VSV pseudotyped viruses (Fig. 1). Additionally, we found that the infectivity titers of VSV-He/80 and VSV-huP2br varied significantly due to a different residue in GP2 (Fig. 2). This residue was critical for membrane fusion activity and determined the pH threshold for membrane fusion (Fig. 3). Our results indicate that the efficiency of BoDV-1-G-mediated cell entry is regulated by its membrane fusion activity and the pH threshold for membrane fusion.

There has been a previous report on the production of VSV-BoDV-1 using the He/80 strain^16^. Interestingly, the infectivity titer of VSV-He/80 generated in our study was significantly lower than that reported previously. Despite the general belief about the high stability of the BoDV-1 genome, various variants have been found within infected cells, which may explain the observed difference in infectivity titer^30^. Analysis of the BoDV-1 strains in the National Center for Biotechnology Information (NCBI) database demonstrated that 113 out of 117 strains, including He/80, have an isoleucine at BoDV-1-G residue 307, whereas the remaining 4 strains, including huP2br, have a methionine, thus suggesting that our findings are not indicative of particular minor variants. Alignment of orthobornavirus G used in this study showed that BoDV-2, PaBV-2, and PaBV-4 possess isoleucine, whereas VSBV-1, CnBV-1, and PaBV-5 possess valine at the position corresponding to BoDV-1-G residue 307 (Fig. 4A). Although the VSV pseudotyped virus with BoDV-2-G exhibited a relatively high infectivity titer, those with PaBV-2 and PaBV-4-G showed lower titers compared to others (Fig. 1A). This suggests that substituting the isoleucine at the position corresponding to BoDV-1-G residue 307 with methionine or valine may increase the infectivity titer.

**Fig. 4.**
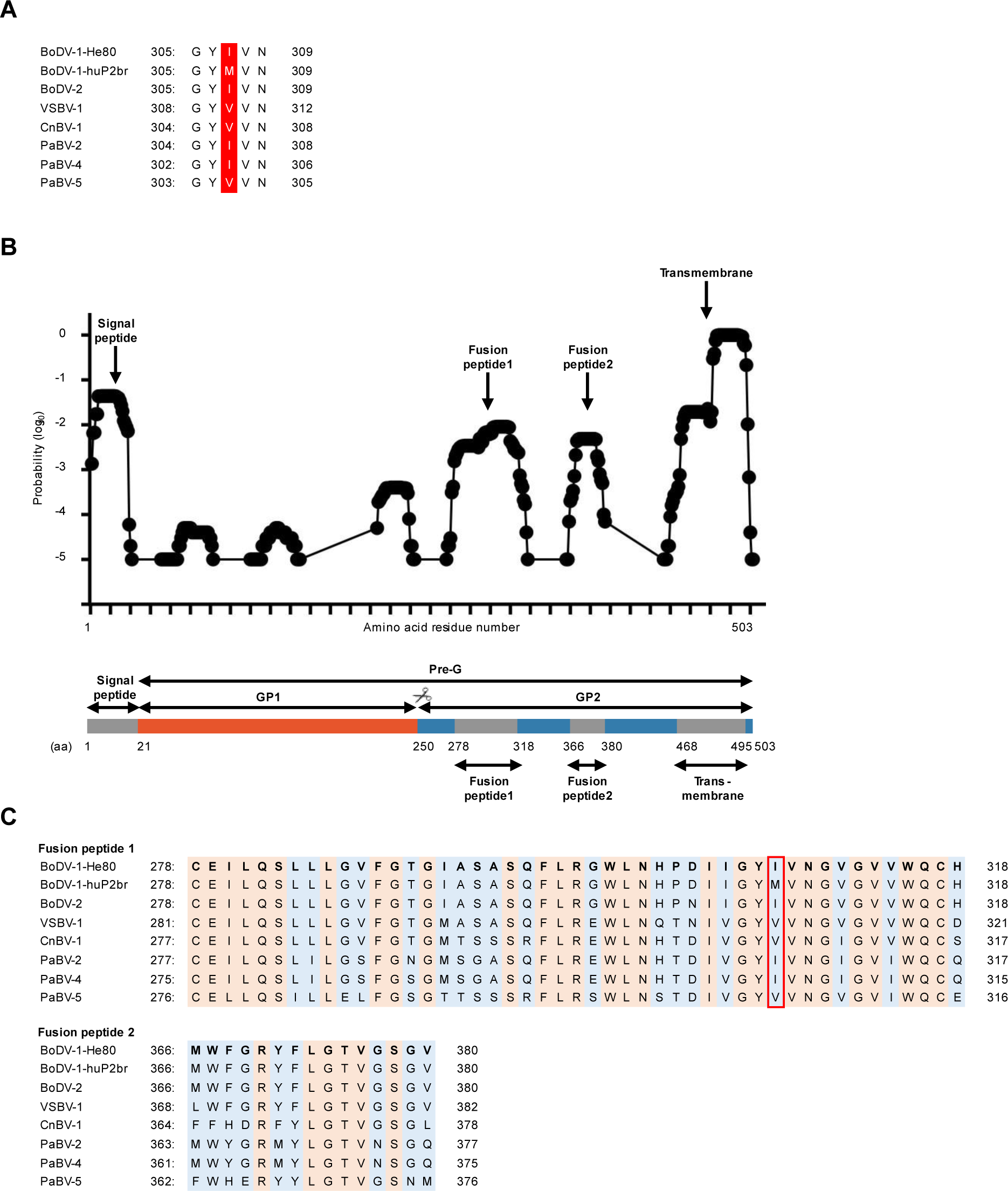
Predicted fusion peptide of BoDV-1. (A) Multiple sequence alignment of residues corresponding to BoDV-1-G 307. The residues corresponding to BoDV-1-G 307 are highlighted in red background. (B) A diagram showing the locations of the signal peptide, fusion peptide, and transmembrane domain predicted by TMHMM. (C) Multiple sequence alignment of the predicted fusion peptides of orthobornavirus used in this study. The conserved residues are highlighted in orange, whereas the nonconserved residues are highlighted in blue. The residues corresponding to BoDV-1-G 307 are surrounded by red line.

VSV buds from the cell surface, incorporating membrane proteins into particles^31^. BoDV-1-G primarily localizes to the ER and Golgi apparatus but also to the plasma membrane^32,33^. Indeed, BoDV-1-G was incorporated into VSV particles (Fig. 3A). Notably, the incorporation of He/80 I307M mutant G into VSV particles was less than that of other G. This mutation reduced the cleavage efficiency of G within the cells (Fig. 2B), thus potentially impairing its transport to the plasma membrane and subsequent incorporation into VSV particles. The mechanism underlying the reduced cleavage efficiency due to this mutation remains unclear, which is mainly due to the absence of structural information on BoDV-1-G. However, it is speculated that this mutation may limit the accessibility of furin to the cleavage site. Conversely, the S56P mutation increased the cleavage efficiency (Fig. 2B), thus suggesting that it may compensate for the reduced cleavage efficiency of He/80 I307M mutant G, thereby enhancing its incorporation into VSV particles and the infectivity titer. Previous reports have shown that mutations in the ER retention signal in envelope proteins, which are predominantly localized to the ER-Golgi apparatus, facilitate their localization to the plasma membrane and enhance their incorporation in VSV particles and infectivity titer^26–28^. BoDV-1-G also contains an ER retention signal within its cytoplasmic tail^33^, and deletion or mutation of this domain may similarly increase the infectivity titer of VSV-BoDV-1.

Viral membrane fusion proteins are categorized into three classes. Some class I fusion proteins, such as the influenza virus hemagglutinin (HA) and coronavirus spike (S) proteins, require cleavage by host proteases to expose a fusion peptide, which is rich in hydrophobic amino acids and essential for membrane fusion. In contrast, most class III fusion proteins, including VSV-G, do not require cleavage by host proteases for membrane fusion. Although a previous computational proteomics analysis classified BoDV-1-G as a class III fusion protein, it needs to be cleaved into GP1 and GP2 by furin for infection^34^. Additionally, the N-terminal domain of GP2 is rich in hydrophobic amino acids, thus suggesting that this domain may function as a fusion peptide, which resembles those in class I fusion proteins^17^. Mutations close to fusion peptides have been reported to alter membrane fusion activity and the pH threshold for fusion in other viral envelope proteins^25^. Since the BoDV-1-G residue 307 affects the cleavage efficiency, this residue may be close to the cleavage site and the GP2-N-terminal region, thus suggesting that this residue affects membrane fusion activity through a similar mechanism. Recently, we showed that the cleavage of BoDV-1-G is crucial for its maturation and glycosylation, implying that this cleavage may not be directly related to membrane fusion^13^. Furthermore, a previous report showed that GP1 alone can facilitate cell entry, which implies that GP1 contains a fusion peptide^16^. This finding is consistent with a previous computational proteomics analysis, although experimental validation has not been performed^34^. Considering that fusion peptides insert into the host membrane, fusion peptides may share properties with transmembrane domains (TMDs). In fact, TMD prediction software has successfully identified fusion peptides in HIV Env, influenza A virus H1N1 HA, and betacoronavirus S proteins^35^. Therefore, we used the TMD prediction algorithm TMHMM to examine the fusion peptides of BoDV-1-G and identified two candidate regions in GP2 (Fig. 4B and C). Mutations within fusion peptides have been reported to alter membrane fusion activity and the pH threshold for fusion in other viral envelope proteins^36,37^. BoDV-1-G residue 307 is included in the predicted fusion peptides, thus suggesting that this residue affects membrane fusion activity through a similar mechanism. Detailed analysis of the BoDV-1-G structure and systematic mutational analysis to identify fusion peptides will provide deep insights into the underlying mechanism.

In other viruses, mutations that alter the pH threshold for membrane fusion affect not only replication and infectivity in cultured cells but also transmission and pathogenicity^29,38,39^. Viruses that infect via the respiratory route are exposed to a mildly acidic extracellular environment in the airways^40^. Therefore, the balance between the facility of membrane fusion in the endosome and stability in the extracellular acidic environment is crucial. Although the exact infection route of BoDV-1 remains unknown, BoDV-1 is presumed to infect via the nasal mucosa and subsequently spread to the central nervous system, thus suggesting that this balance may also be important for BoDV-1 infection^41,42^. In addition, mutations that increase membrane fusion activity have been reported to increase the pathogenicity of neurotropic viruses upon intracerebral inoculation^43^. Further studies are required to determine whether variations in membrane fusion activity and the pH threshold for membrane fusion of BoDV-1-G affect infectivity, transmission, and pathogenicity.

In this study, we successfully developed an efficient method for producing VSV pseudotyped viruses with orthobornavirus G. Furthermore, we demonstrated that both the membrane fusion activity and the pH threshold for membrane fusion play crucial roles in determining the infectivity titer. This study provides valuable insights into the cell entry mechanism of orthobornaviruses and paves the way for the development of drugs that inhibit cell entry and diagnostic systems. Additionally, our findings contribute to elucidating the properties of orthobornavirus G and identifying its functional domain.

## Funding information

Japan Society for the Promotion of Science (JSPS) KAKENHI, Grant Numbers: JP23K14082 (HM), JP20H05682 (KT), JP21K19909 (KT)

Japan Agency for Medical Research and Development (AMED), Grant Number: JP23bm1223017 (KT) Kaketsuken Research Grant (KT)

The Joint Usage/Research Center Program on Institute for Life and Medical Sciences, Kyoto University (KT)

## Author contributions

Conceptualization, methodology, investigation, and writing: Yusa Akiba.

Conceptualization, methodology, writing, supervision, and funding acquisition: Hiromichi Matsugo. Methodology and resources: Takehiro Kanda.

Methodology and resources: Madoka Sakai. Methodology and resources: Akiko Makino.

Supervision, writing, and funding acquisition: Keizo Tomonaga.

## Abbreviations

ABV: avian bornavirus
BoDV-1: Borna disease virus 1
CnBV-1: canary bornavirus 1
DMEM: Dulbecco’s modified Eagle’s medium
ER: endoplasmic reticulum
FCS: fetal calf serum
G: glycoprotein
GFP: green fluorescence protein
HA: hemagglutinin
L: RNA-dependent RNA polymerase
M: matrix protein
N: nucleoprotein
P: phosphoprotein
PaBV: parrot bornavirus
PBS: phosphate-buffered saline
PDD: proventricular dilatation disease
S: spike
TMDs: transmembrane domains
vRNPs: viral ribonucleoproteins
VSBV-1: variegated squirrel bornavirus 1
VSV: vesicular stomatitis virus

## Acknowledgments

This study was supported in part by JSPS KAKENHI grants JP23K14082 (HM), JP20H05682 (KT), JP21K19909 (KT); AMED under Grant Number, JP23bm1223017 (KT); Kaketsuken Research Grant (KT); the Joint Usage/Research Center Program on Institute for Life and Medical Sciences, Kyoto University (KT).

